# Scan Cluster: A versatile database-independent prediction tool for multi-genome identification of homologous gene clusters

**DOI:** 10.64898/2026.04.29.721675

**Authors:** Ezequiel G. Mogro, Abril L. Pagnutti, Gonzalo Zapata, Mauricio J. Lozano

## Abstract

The exploration of colocalized gene sets, such as Biosynthetic Gene Clusters (BGCs) and symbiotic islands, is fundamental in modern genome mining. However, many existing prediction tools rely heavily on curated databases or predefined rules, inherently biasing detection toward known clusters. To address these limitations, we introduce Scan Cluster, a robust and flexible Python-based bioinformatic tool designed to identify user-defined, conserved gene clusters across diverse genomes without database-driven constraints. Scan Cluster leverages BLAST or HMMER to detect homologous proteins and evaluates their colocalization, accommodating complex evolutionary events such as orientation inversions, the insertion of alien genes, gene deletions, and the integration of insertion sequences. Beyond identification, the software performs progressive multiple cluster alignments and generates distance trees to assess cluster similarity, producing outputs ready for visualization in iTOL and Clinker. We validated Scan Cluster against standard tools like antiSMASH and DeepBGC, demonstrating high accuracy in delineating complete cluster boundaries. Its utility was further confirmed through the analysis of the *nos* and complex *nod-nif-fix* symbiotic gene clusters in rhizobia strains, successfully tracking genetic decay and grouping diverse architectures. Scan Cluster provides an accessible, low-resource framework to explore the evolutionary and functional dynamics of novel genetic clusters.

## Introduction

Biosynthetic Gene Clusters (BGCs) are spatially co-localized DNA segments encoding proteins involved in the biosynthesis of diverse natural products, commonly referred as specialized or secondary metabolites (Chavali & Rhee, 2017; Liu et al., 2022). These molecules represent an invaluable reservoir of bioactive compounds with profound implications across medicine and agriculture (Cavas & Kirkiz, 2022; Hannigan et al., 2019). Secondary metabolites serve as crucial sources for pharmaceuticals, including antibiotics and anticancer agents, and play vital roles in ecological interactions such as defense and signaling (Chavali & Rhee, 2017). BGCs can be horizontally transferred and evolve rapidly, usually showing exceptionally high rates of insertions, deletions, duplications and rearrangements (Medema et al., 2014). They facilitate the synchronized regulation of gene expression and the inheritance of metabolic pathway blocks —sub-clusters— that can be further recombined, allowing bacteria to rapidly acquire complex traits. Microorganisms, particularly bacteria and fungi, are the main subject of research for producing these compounds, often synthesizing them via complex biosynthetic pathways encoded within BGCs (Anand et al., 2010; Chavali & Rhee, 2017; Liu et al., 2022).

While biosynthetic clusters are essential targets of study, a wide variety of relevant cellular functions are also encoded within colocalized genes. In this context, genes involved in symbiotic functions provide a particularly relevant example. For instance, in rhizobia (soil bacteria that fix nitrogen when associated with legume plants), the genes necessary for nodulation (*nod*) and nitrogen fixation (*nif* and *fix*) are frequently arranged in gene clusters (Cafiero et al., 2023; Iismaa et al., 1989; Nichio et al., 2025). Nodulation genes, in particular, are characterized by high rates of horizontal gene transfer (HGT) between different rhizobial lineages (Epstein & Tiffin, 2021; Sugawara et al., 2013), sometimes flanked by insertion sequences that facilitate their mobility (Epstein & Tiffin, 2021; Teulet et al., 2020).

The advent of high-throughput genome sequencing has revolutionized natural product discovery, providing a new perspective through a computational approach to explore microbial genomes for novel BGCs. This shift in focus has led to the development of numerous bioinformatic tools designed to identify and characterize these genetic clusters. For example, antiSMASH is recognized for its comprehensive pipeline and rule-based detection of known BGC types using profile HMMs (Blin et al., 2025), and is widely used to identify common cluster types. Machine learning methods were incorporated in BGC prediction software like ClusterFinder and DeepBGC. ClusterFinder uses a two states (BGC and non-BGC) hidden Markov model trained on genomic pFAM domains that allows the detection of both known and putative novel cluster types (Cimermancic et al., 2014). However, it was reported to predict shorter BGCs and to have many false positive predictions (Hannigan et al., 2019). DeepBGC employs a Bidirectional Long Short-Term Memory (BiLSTM) Recurrent Neural Network that incorporates contextual gene information improving accuracy and enabling the detection of novel BGC classes (Hannigan et al., 2019).

Additional tools complement these approaches with more specialized capabilities. PRISM integrates genome mining with cheminformatics to predict both BGCs and the chemical structures of their associated natural products (Skinnider et al., 2020). BAGEL focuses specifically on the identification of bacteriocin gene clusters using curated databases and motif-based approaches (van Heel et al., 2018). cblaster enables the identification of homologous gene clusters across local and remote databases by leveraging BLAST-based searches and genomic colocalization (Gilchrist et al., 2021). During the last few years BGC prediction methods based on the transformer-based language model architecture or protein language model (pLM) embeddings have been reported showing remarkable results (Kang et al., 2025; Lai et al., 2025; Rios-Martinez et al., 2022).

However, despite the wide range of available tools, important limitations remain. Several earlier methods and associated databases are now outdated, superseded, or no longer actively maintained, reducing their applicability in current genome mining workflows. Moreover, many widely used tools, including PRISM, BAGEL, and cblaster, rely heavily on curated databases, sequence homology, or predefined domain profiles for BGC identification, which inherently biases detection toward previously characterized biosynthetic systems. While approaches such as ClusterFinder reduce direct database dependency through probabilistic modeling, they still inherit biases from their training data. Regarding the newer methods utilizing pLMs, the main limitation for the common user is the requirement of GPUs. Altogether, these limitations highlight the need for approaches capable of accurately identifying gene clusters beyond database-driven constraints and with low computational resources.

In this context, the continuous discovery of new natural products (Yushchuk et al., 2025) and the increasing volume of genomic data necessitate the development of more versatile tools capable of identifying homologous gene clusters across diverse organisms and facilitating the exploration of novel cluster architectures.

Here we introduce Scan Cluster, a bioinformatic tool designed to address these limitations by providing a robust and flexible framework for identifying homologous gene clusters in bacterial genomes, with potential applicability to eukaryotes and viruses. Scan Cluster distinguishes itself by offering greater flexibility and customization, enabling the identification of user-defined, syntenically conserved gene sets. Further, multiple progressive cluster alignment and cluster distance trees are generated to assess cluster similarity. The software can process hundreds of bacterial genomes with low end hardware requirements and is also available as a Google Colaboratory notebook. Scan Cluster thus aims to streamline the laborious process of gene cluster searching and annotation, offering researchers an accessible and powerful tool to gain deeper insights into the evolutionary and functional aspects of these genetic elements.

## Materials and methods

### Scan cluster algorithm

Scan Cluster is a program written in python using the Biopython (Cock et al., 2009) and pandas (Reback et al., 2020) libraries; and based on homologous protein searches using either Blastp (Altschul et al., 1990) or hmmsearch from the HMMER suit (Eddy, 2011).

The program has three main running modes that differ in the first steps of the algorithm which is responsible for the search of homologous proteins.

The default running mode starts from a query cluster that can be provided as a genbank format file, or as a reference genome file in genbank format along with the replicon accession number and the cluster start/end coordinates. In the latter case, the query cluster is extracted as a genbank file from the reference genome. The following step uses either local or remote Blastp to search for homologs of each protein in the query cluster. Those homologs are then filtered by specified coverage and E-value thresholds, and aligned with mafft (Katoh & Standley, 2013) using the L-INS-I mode. The Blastp search can be done using the *-remote* option using any of the NCBI protein databases, or in local mode which uses either a Blast formatted database provided by the user or database generated by the program using all the target genomes. From the resulting multiple sequence alignment of the query cluster proteins a HMM is built with hmmbuild. As a result HMM profiles for each of the proteins of the query cluster are obtained and saved in the results folder.

Alternatively, scan_cluster can search for clusters containing a predefined set of HMM, included in a user defined folder. Finally, it can search for homologs of the proteins in the query cluster using Blastp instead of hmmsearch. In that case HMMs are not used, being the homolog protein search performed in a less sensitive sequence-sequence mode.

Next, the hmmsearch (or blast) result tables are parsed and analyzed to identify colocalized protein hits —the proteins are enumerated and their order is analyzed— to define clusters following a set of user customizable rules. A cluster is defined if: a) there are multiple query protein hits (-m, --min_target_prots, default value of 3) in the same region of the target genome; b) the homologs of the query proteins are not separated in the target genome by more than -n/--n_prots_between proteins that are not a part of the query cluster; c) there are less than -M/--max_alien_prots proteins in the target cluster that are not present in the query cluster (alien proteins); and d) the fraction of the query proteins found in the target cluster is greater than --min_cluster_coverage (Default value of 0.5, meaning that the program will keep clusters containing homologs of at least half of the query proteins). If the program is run in the HMM mode, with a large collection of HMMs, this argument should be carefully tuned, in particular if clusters are expected to contain only a fraction of the HMM collection.

The final step involves the generation of a scoring system for the pairwise cluster alignment by dynamic programming, and the progressive multiple cluster alignment. A scoring system for the matched proteins is generated by calculating the percent identity of all homologous protein pairs. For the proteins that are part of the target clusters but were not in the query cluster, an all vs all Blastp is performed to search for homologs within the clusters. Those proteins are aligned using mafft L-INS-I mode (if more than 100 proteins need to be aligned Scan cluster will switch to --fast mode), and pairwise percent identities computed. For aligned genes with different orientations, the match score is halved. Fixed (but customizable) mismatch and gap scores are used that favor the gap introduction over the mismatches in order to get the best alignment of the homolog proteins.

All clusters are aligned to the reference cluster and reoriented to maximize the alignment score, so that most of the genes are in the same direction as the query cluster. Next, all the pairwise cluster alignments are computed, a distance matrix is generated and used to build a UPGMA distance guide tree using the biopython Phylo library. Finally, the progressive alignment is computed following the guide tree, aligning the closest clusters first, and using the sum of scores method to align sub cluster alignments.

As a result, scan_cluster generates multiple files including the identified clusters, a distance tree of the clusters, and an annotation file to use with Itol tree visualizations (Letunic & Bork, 2019). The clusters are also exported in genbank format to perform an independent cluster analysis with Clinker software (Gilchrist & Chooi, 2021).

### Data Acquisition and HMM Generation

The genomes used in this study were retrieved from the NCBI Genome database, prioritizing RefSeq assemblies. Custom Hidden Markov Model (HMM) profiles were obtained as follows: Amino acid sequences for each gene in each of the clusters were obtained from UniProt. These searches were based on the most common gene annotation, exclusively selecting sequences from the manually curated Swiss-Prot database. The protein sequences were then aligned using MUSCLE, and the resulting alignments were converted to Stockholm format to generate the final HMM profiles using *hmmbuild*.

### Cluster identification and visualization

The analysis of biosynthetic gene clusters (BGCs) was performed using antiSMASH version 8.0 and DeepBGC version 0.1.31, both with default parameters. Scan cluster was used with default options unless stated otherwise (see Software settings and parameters in Supplementary Information). For the generation of the cluster figures clinker (v0.0.31) was used with the identified clusters extracted in genbank format from the corresponding genomes. The Interactive Tree of Life server (iTOL v7, https://itol.embl.de/) was used to generate the Scan cluster annotated tree figure using the generated tree files and Itol annotation files.

## Results and Discussion

### Searching and aligning clusters using Scan cluster

Here we introduce Scan cluster (scan_cluster.py), a program implemented in the python language, designed to search for any user-defined set of colocalized genes in multiple target genomes. Scan cluster goes further, generating a multiple cluster alignment (MCA) and providing a way to assess cluster similarity.

Briefly, given a query cluster, Scan cluster searches for homologs of all the proteins in the cluster using either HMM profiles or the aminoacid sequence. The results are processed to identify colocalized proteins and clusters are identified using a set of simple and fully customizable rules. Next, a scoring system is generated based on homolog protein percent identities and an all-vs-all pairwise dynamic programming global cluster alignment using the Needleman-Wunsch algorithm is performed. The pairwise alignment scores are used to generate a distance matrix that is subsequently used to generate a guide tree for the progressive multiple cluster alignment.

The program is freely available under MIT licence in github (https://github.com/maurijlozano/scan_cluster) and zenodo (https://doi.org/10.5281/zenodo.15195352) and can be run locally as a command line program or jupyter notebook. Additionally, to make it more accessible to researchers with less bioinformatic skills, scan_cluster is also available as a Google Colaboratory notebook (https://colab.research.google.com/github/maurijlozano/scan_cluster/blob/main/scan_cluster.ipynb), providing a clear form like user interface. This notebook also incorporates a module to download the query cluster (reference genome plus replicon ID and cluster genomic coordinates), and target genomes from NCBI using the datasets tool (O’Leary et al., 2024).

### Scan cluster algorithm validation and feature demonstration

To rigorously evaluate the performance of Scan Cluster, we conducted a comparative analysis against two of the most widely recognized and state-of-the-art bioinformatics tools for Biosynthetic Gene Cluster (BGC) prediction: antiSMASH 8.0 (Blin et al., 2025) and DeepBGC (Hannigan et al., 2019). Our goal is not to conduct an exhaustive benchmark against these specialized tools, which are optimized for identifying a large number of established Biosynthetic Gene Clusters (BGCs). Instead, we aim to demonstrate the strong performance of Scan cluster when using an appropriate set of cluster definition arguments, tailored by the researcher’s knowledge of the specific genetic cluster.

This initial assessment was performed on three *E. coli* genomes (K-12, Nissle 1917, VR50; accession numbers for these strains are provided in Table S1) through the identification of the well-characterized gene cluster for enterobactin synthesis (Cavas & Kirkiz, 2022; Gehring et al., 1997; Nahlik et al., 1987). That Scan cluster perfectly matched the core predictions of antiSMASH (Figure 2) is particularly significant, as it is considered a global reference standard due to its exhaustive manual curation and consolidated heuristic rules. In contrast, the prediction made by DeepBGC yielded a significantly smaller genome segment for this cluster, where regulatory, transport-related, and various other auxiliary genes that contribute to the overall biological process were not detected.

**Figure 1.**
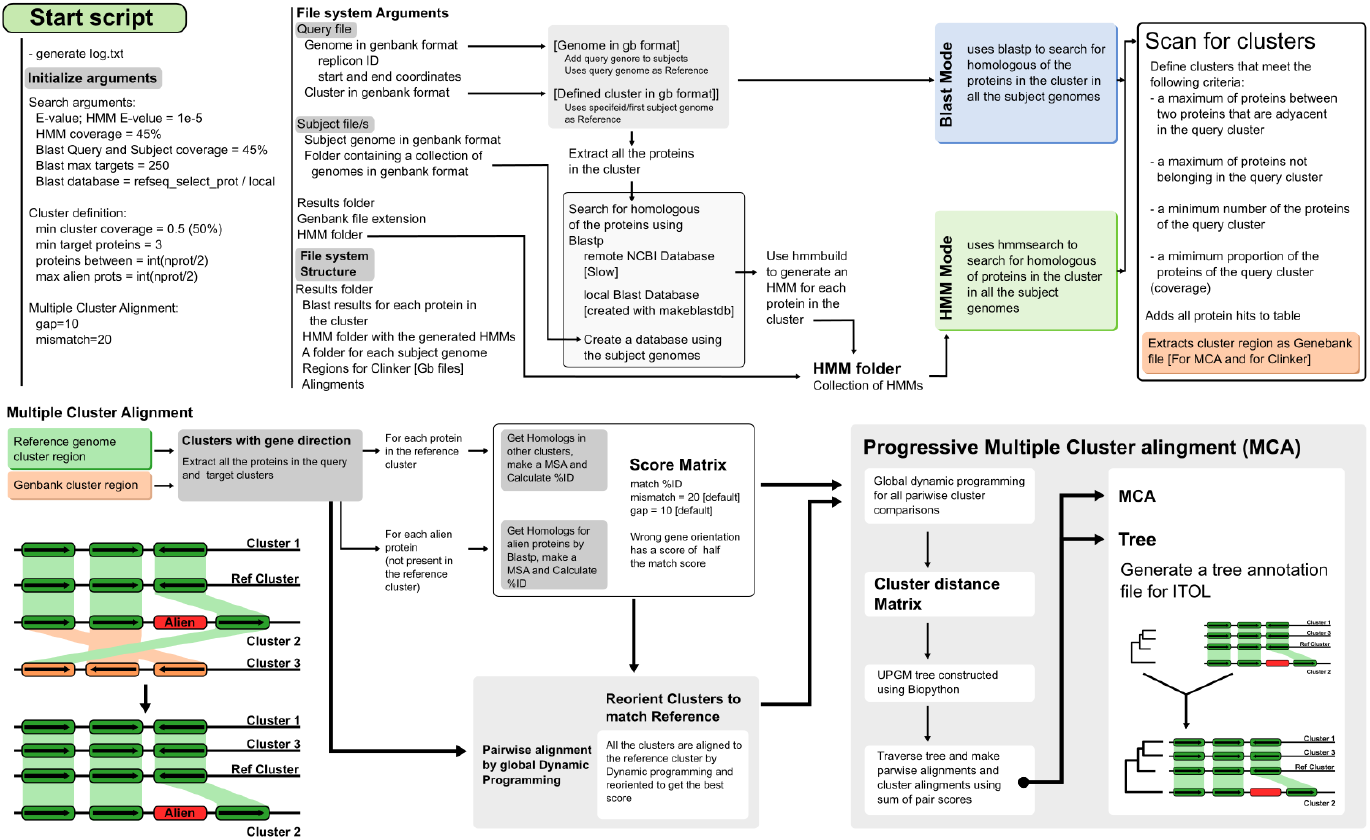
Scan cluster algorithm. A summary of Scan cluster algorithm detailing the search, cluster definition and MCA arguments; the different required inputs and the corresponding running modes.

**Figure 2.**
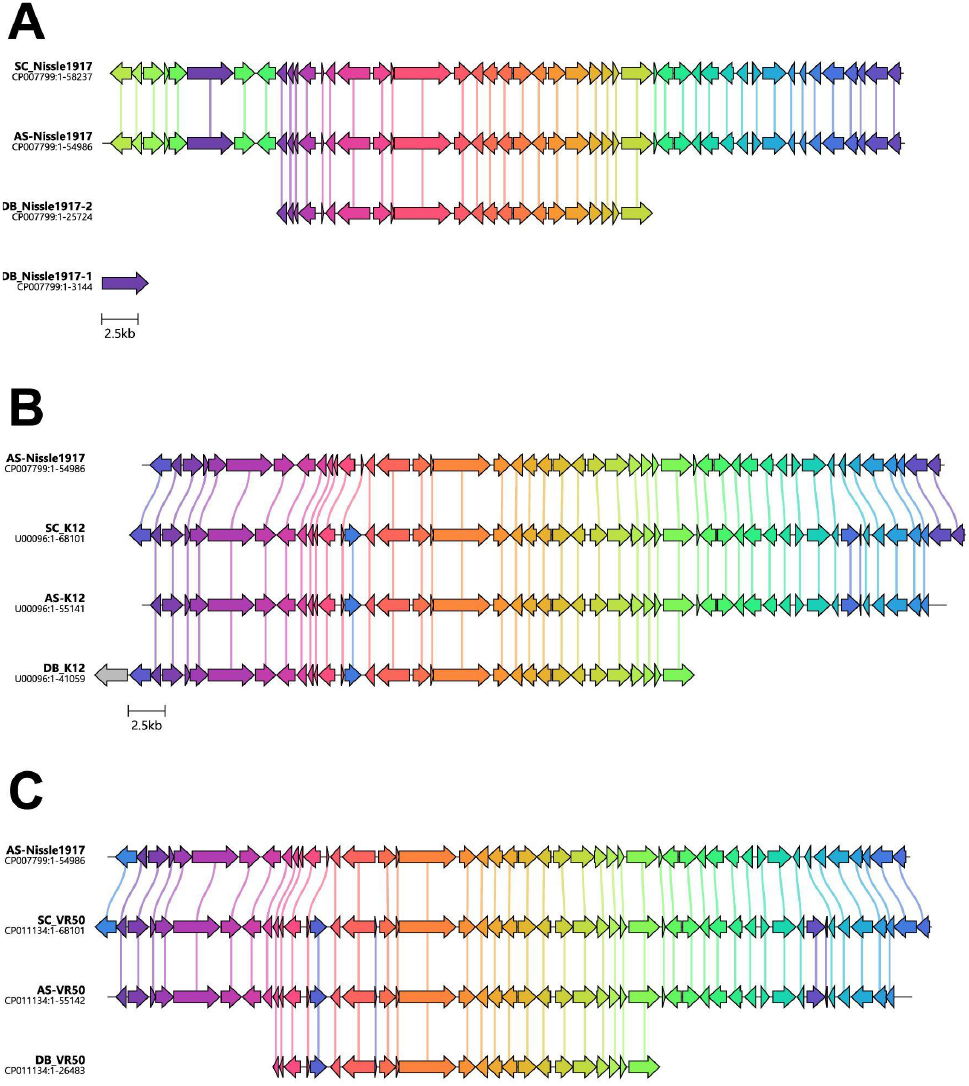
Comparison of the enterobactin gene cluster in *E. coli* Nissle 1917 **(A)**, K-12 **(B)**, and VR50 **(C)** strains predicted by Scan Cluster (SC), antiSMASH (AS), and DeepBGC (DB). The antiSMASH-predicted enterobactin cluster from strain Nissle 1917 was used as the reference for all Scan Cluster runs. The comparative visualization among the different strains was generated with Clinker using the cluster regions exported by Scan Cluster. For DeepBGC, the region was obtained by extracting the location specified in the results from the input GenBank file. For antiSMASH, the cluster was obtained from the output files. The vertical connecting lines establish sequence homology, indicating a protein identity level greater than 30% between the corresponding genes.

This finding indicates a limitation in DeepBGC’s ability to capture the complete genetic landscape of a BGC. Because its boundary detection relies on a supervised learning approach, the model is inherently constrained by the negative examples present in its training data (Rios-Martinez et al., 2022). This reliance makes it prone to false negatives at the sequence edges, which can prematurely truncate the predicted cluster and exclude relevant peripheral genes. In contrast, our results show that Scan Cluster can effectively delineate the full extent of these clusters without depending on database curation.

In addition to searching for BGCs, Scan cluster can be used for the identification of any kind of co-localized protein coding sequences using as input either predefined HMMs for each of the proteins in the cluster, or by passing the cluster as a genbank file or genome file plus cluster start and end coordinates. This Scan Cluster’s unique feature was evaluated through the identification of *nos* gene clusters in *Sinorhizobium meliloti* (Figure 3). The *nos* cluster (*nosRZDFYLX* genes) plays a key ecological role by encoding nitrous oxide reductase, an enzyme that reduces the greenhouse gas nitrous oxide (N_2_O) to nitrogen gas (N_2_), helping to lower emissions in agriculture. The *nos* gene cluster in *S. melilot* was previously reported to be present in four laboratory strains (1021, BL255C, RM41 and GR4), and absent in certain commercial isolates (B399 and B401) as a consequence of substantial genetic decay, characterized by extensive structural deletions and mutational events associated with rhizobial domestication (Brambilla et al., 2018). Scan Cluster search using as input custom made HMMs for the 7 *nos* proteins (Nos X, L, Y, F, D, Z and R; Figure 3) successfully identified the cluster in *S. meliloti* 1021, BL225C, RM41 and GR4 but not in B399 and B401 strains. These results demonstrate the utility of Scan Cluster for assessing strain quality by accurately detecting the retention or loss of metabolic clusters of interest.

**Figure 3.**
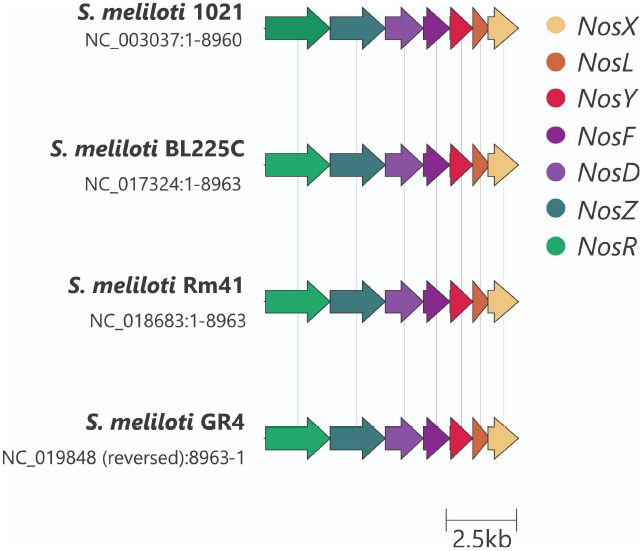
*nos* gene cluster in *S. meliloti* laboratory strains. The comparative visualization among the different strains was generated using the Clinker using the cluster regions exported by Scan Cluster. The *nos* cluster was not found in *S. meliloti* B399 and B401 strains; hence, their absence in Scan cluster results. The vertical connecting lines establish sequence homology, indicating a protein identity level greater than 30% between the corresponding genes.

Scan cluster also performs cluster comparison by dynamic programming (see Materials and Methods) and progressive multiple cluster alignment. From the output files, a cluster distance tree and an annotation file for iTol (Letunic & Bork, 2019) can be obtained, which greatly simplify the synteny and cluster comparison. To evaluate the grouping of clusters by similarity we analyzed the gene architecture of the *nod, nif*, and *fix* gene clusters, where concordant results with those reported in the literature by De Meyer et al. (2016) were found in rhizobia strains of different species (Figure 4). This is a difficult task that will normally yield fragmented genomic regions corresponding to each cluster, since *nod-nif-fix* clusters were observed exhibiting diverse architectures, notably including a high number of alien genes and inversions of specific regions within the clusters. After running Scan Cluster with default parameters, it was observed that many clusters were rejected but annotated in the discarded_clusters output file. Initial analysis with Scan Cluster yielded results for only a small proportion of the genomes, as many clusters were detected in fragments and thus discarded (low cluster coverage). While expanding the maximum allowed gap between consecutive genes (-n = 20) successfully mitigated this fragmentation, it simultaneously led to the prediction of artificially elongated clusters. To counteract this and establish stricter cluster boundaries, the coverage thresholds for the BLAST homology searches (--Blast_scov and --Blast_qcov) were raised to 65%. In figure 4, we show that given the appropriate arguments (based on minimal knowledge of the cluster architecture) Scan cluster could find the query genomic region in all the target genomes. This makes Scan cluster a unique tool to search for colocalized genes, even in highly dynamic genomic regions.

**Figure 4.**
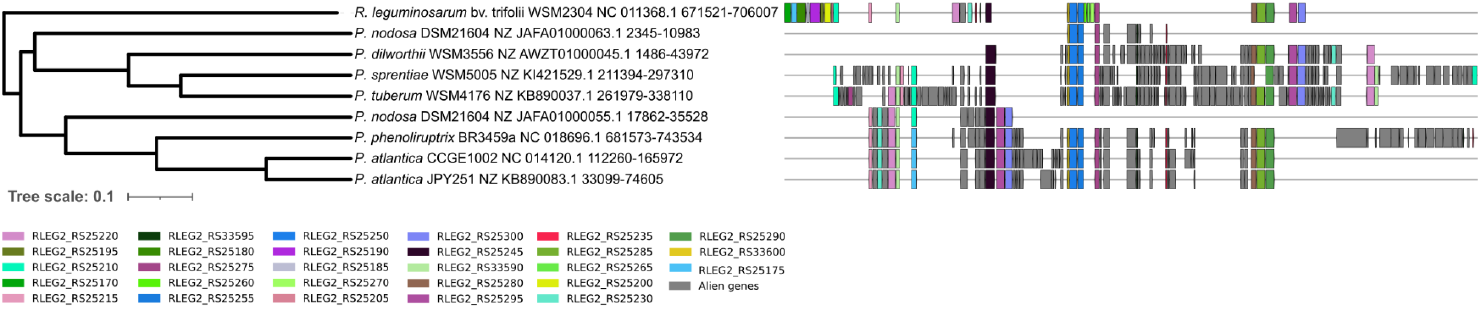
Prediction of the *nod, nif*, and *fix* cluster structures performed by Scan Cluster using HMM profile searches. Graphical representation of the cluster distance tree obtained via iTOL. The annotation file generated by Scan cluster was used to annotate the tree, showing homologs of the query genes in different colors. Blocks in the same position correspond to aligned proteins. Gray colored blocks correspond to alien proteins. White spaces correspond to insertions or deletions.

### Comparative analysis of symbiotic genes in *Sinorhizobium meliloti*

Scan cluster’s capability to identify homolog clusters was also assessed by conducting a comparative genomic analysis of the symbiotic genes (*nod, nif*, and *fix*) in *Sinorhizobium meliloti*, a model symbiotic nitrogen fixing bacteria. In this case, the reference genomic region from *S. meliloti* 2011 was passed as query —reference genome file, replicon accession number, and the corresponding start and end coordinates— and the search was done using the profile HMMs generated by Scan cluster (Figure 5; see Material and Methods section for detailed running options).

**Figure 5.**
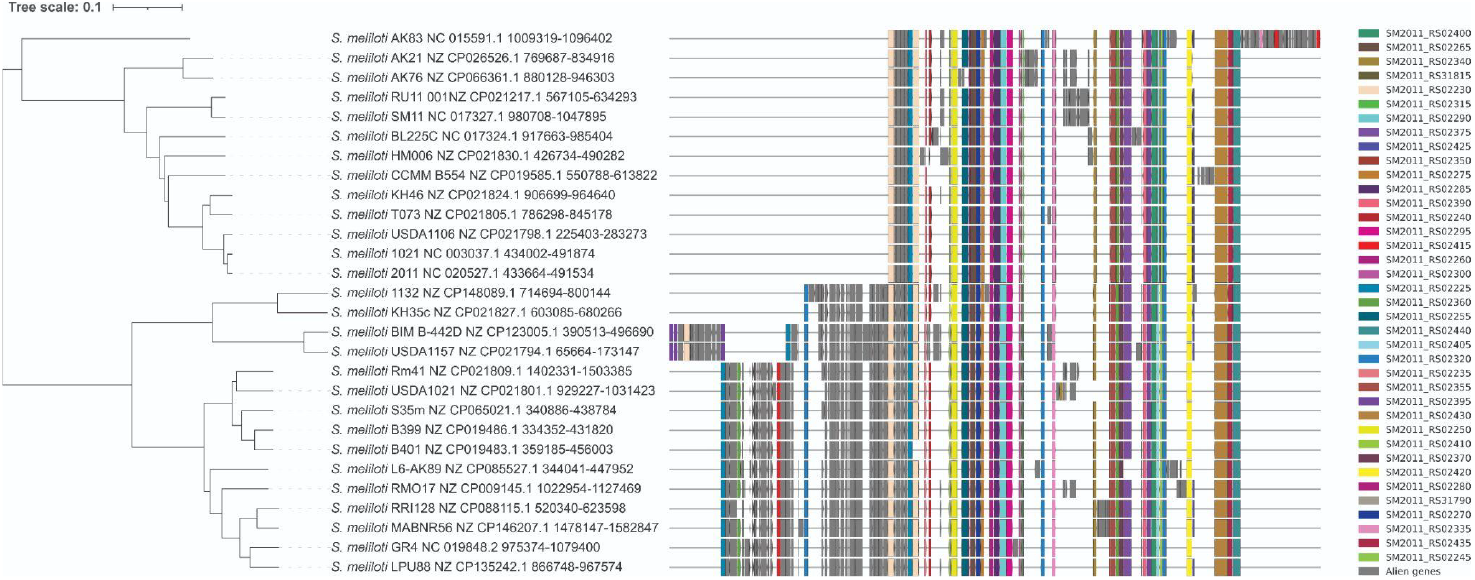
Prediction of *nod, nif*, and *fix* gene cluster structures in pSymA megaplasmids of *S. meliloti* strains. The analysis included all complete *S. meliloti* genomes from the RefSeq database and was performed by Scan Cluster, using the cluster present in strain 2011 as the query. The graphical representation of the cluster distance tree was generated via iTOL and annotated using the Scan Cluster iTOL annotation output file. Different colors indicate homologs of the query genes. Blocks in the same vertical position correspond to aligned proteins, gray blocks represent alien proteins, and white spaces indicate insertions or deletions.

When contrasting the Scan Cluster results with those obtained by OrthoFinder (Table S5, Emms & Kelly, 2019), a high overall accuracy was observed; however, OrthoFinder detected 4 additional genes that Scan Cluster omitted. These elements are annotated as pseudogenes in the reference genome, and thus are intentionally discarded by the Scan Cluster algorithm during its processing.

## Conclusions

The continuous discovery of natural products and the exponential growth of genomic data require versatile bioinformatic tools that overcome the biases of strictly curated databases. We address this need by providing a flexible, robust framework for the identification of user-defined, syntenically conserved gene sets across diverse genomes. Scan Cluster can identify various types of gene clusters using either BLAST or HMMER, accommodates complex evolutionary events (inversions, insertions, deletions), and operates with low computational resources, even being available as a Google Colaboratory notebook. Its high flexibility allows Scan cluster to identify almost any kind of colocalized gene sets, provided the researcher has minimal architectural knowledge of the target region.

We evaluated Scan Cluster’s performance in a set of different tasks. First, we show that Scan cluster achieves a high level of concordance with established reference standards like antiSMASH, accurately defining the full extent of gene clusters without prematurely truncating peripheral genes, a limitation observed in machine-learning models such as DeepBGC. Although newer artificial intelligence methods based on pLM embeddings or other transformer-based architectures outperform DeepBGC (Kang et al., 2025; Lai et al., 2025; Rios-Martinez et al., 2022) they require higher-end computers with GPUs, which are not always available for the user.

Second, we used Scan cluster to search for the *nos* cluster across various *Sinorhizobium meliloti* strains. The successful identification highlights its utility in tracking genetic decay, assessing strain quality, and guiding the bioprospecting of high-quality strains for the development of bio-inputs. Third, by analyzing complex symbiotic gene clusters (*nod-nif-fix*), we showed that the algorithm is effective at grouping clusters by similarity, accounting for diverse architectures characterized by gene inversions and the presence of alien genes.

Overall, beyond identification, Scan Cluster optimizes the workflow of gene cluster annotation and comparison. By integrating progressive multiple cluster alignments and generating outputs directly compatible with visualization tools such as iTOL and Clinker, it greatly simplifies synteny analysis. Coupled with its low hardware requirements and user-friendly interface via a Google Colaboratory notebook, Scan Cluster stands as a practical, accessible resource for researchers investigating the structural and adaptive dynamics of genetic clusters.

