## Supplementary information for "Scan Cluster: A versatile database-independent prediction tool for multi-genome identification of homologous gene clusters"

### Dataset of analyzed genomes

The genomes used in this study were retrieved from the NCBI Genome database, prioritizing RefSeq assemblies (Tables S1-S4). To construct the Hidden Markov Model (HMM) profiles, amino acid sequences for each gene in the cluster were obtained from UniProt. These searches were based on the most common gene annotation, exclusively selecting sequences from the manually curated Swiss-Prot database. The protein sequences were then aligned using MUSCLE, and the resulting alignments were converted to Stockholm format to generate the final HMM profiles using *hmmbuild*.

### Software settings and parameters

The following commands were used to compare the results from antiSMASH, DeepBGC, and Scan Cluster. The antiSMASH output for strain Nissle 1917 was used as the query cluster for the Scan Cluster analysis. Given the high conservation of the cluster architecture, the maximum number of proteins between was set to 3 (-n 3).

```
./scan_cluster.py -q ./antismash8_gb/entero_nissle1917.gb -F ./gb_ecoli --Blast_DB nr -n 3
```

The analysis was performed using antiSMASH v8.0.1 and DeepBGC v0.1.31 with default settings.

---

To search for *nos* gene clusters in laboratory and commercial rhizobial strains a custom profile HMM was built from representative proteins selected from Swiss-Prot.

```
./scan_cluster.py -F "./Sme-nos_gb" -f ./nos_hmm/
```

---

To search for *nod*, *nif*, and *fix* clusters in rhizobia, the corresponding cluster from *R. leguminosarum* bv. *trifolii* WSM2304 (accession NC_011368.1, region 671,000-706,300) was used as the query. For this analysis, default parameters were modified to --Blast_DB nr --Blast_qcov, -65 --Blast_scov 65, and -n 20

```
./scan_cluster.py -Q ./gb_burkho/Rhizobium_leguminosarum_bv._trifolii_WSM2304.gb -R NC_011368.1 -s 671000 -e 706300 -F ./gb_burkho/ --Blast_DB nr --Blast_qcov -65 --Blast_scov 65 -n 20
```

---

Search for *nod*, *nif*, and *fix* gene clusters in complete RefSeq genomes of *Sinorhizobium meliloti* strains was performed using the corresponding cluster from *S. meliloti* 2011 (accession NC_020527.1, region 438,249-491,559) as a reference.

```
./scan_cluster.py -Q ./Sme_plasmids_genref/2011__NC_020527.1.gb -R NC_020527.1 -s 438249 -e 491559 -F ./Sme_plasmids_genref --Generate_local_db
```

**Table S1.** Genomes analyzed using antiSMASH, DeepBGC, and Scan Cluster to search for enterobactin biosynthesis genes.

| Strain | Accession Number | Assembly Accession | Reference |
|---|---|---|---|
| *Escherichia coli* str. K-12 substr. MG1655 | U00096.3 | GCA_000005845.2 | Blattner et al., 1997 |
| *Escherichia coli* Nissle 1917 | CP007799.1 | GCA_000714595.1 | Reister et al., 2014 |
| *Escherichia coli* VR50 | CP011134.1 | GCA_000968515.1 | Beatson et al., 2015 |

**Table S2**. Genomes used for the identification of *nif/fix/nod* gene clusters across different rhizobial species.

| Strain | Accession Number | Assembly Accession | Reference |
|---|---|---|---|
| *Paraburkholderia atlantica* CCGE1002 | NC_014117.1 | GCF_000092885.1 | Ormeño-Orrillo et al., 2012 |
| *Paraburkholderia atlantica* JPY251 | NZ_KB890042.1 | GCF_000372985.1 | |
| *Paraburkholderia dilworthii* WSM3556 | NZ_AWZT01000001.1 | GCF_000472525.1 | De Meyer et al., 2015 |
| *Paraburkholderia nodosa* DSM 21604 | NZ_JAFA01000001.1 | GCF_000519185.1 | |
| *Paraburkholderia phenoliruptrix* BR3459a | NC_018695.1 | GCF_000300095.1 | de Oliveira Cunha et al., 2012 |
| *Paraburkholderia sprentiae* WSM5005 | NZ_KI421529.1 | GCF_000473465.1 | |
| *Paraburkholderia tuberum* WSM4176 | NZ_KB890029.1 | GCF_000372945.1 | |
| *Rhizobium leguminosarum* bv. *trifolii* WSM2304 | NC_011369.1 | GCF_000021345.1 | Reeve et al., 2010 |

**Table S3.** Genomes used for the identification of *nos* genes in laboratory and commercial rhizobial strains.

| Strain | Accession Number | Assembly Accession | Reference |
|---|---|---|---|
| *Sinorhizobium meliloti* 1021 | NC_003047.1 | GCF_000006965.1 | Capela et al., 2001 |
| *Sinorhizobium meliloti* B399 | NZ_CP019488.1 | GCF_002302375.1 | |
| *Sinorhizobium meliloti* B401 | NZ_CP019485.1 | GCF_002302355.1 | |
| *Sinorhizobium meliloti* BL225C | NC_017322.1 | GCF_000147775.2 | Galardini et al., 2013 |
| *Sinorhizobium meliloti* GR4 | NC_019845.2 | GCF_000320385.2 | |
| *Sinorhizobium meliloti* Rm41 | NC_018700.1 | GCF_000304415.1 | |
| *Sinorhizobium meliloti* SM11 | NC_017325.1 | GCF_000218265.1 | Schneiker-Bekel et al., 2011 |

**Table S4.** pSymA megaplasmids used for the identification of *nif*/*fix*/*nod* gene clusters in different *Sinorhizobium meliloti* strains. All complete *S. meliloti* genomes from the RefSeq database were selected for this analysis.

| Strain | Accession Number | Assembly Accession | Reference |
|---|---|---|---|
| *Sinorhizobium meliloti* 1021 | NC_003037.1 | GCF_000006965.1 | Barnett et al., 2001 |
| *Sinorhizobium meliloti* 1132 | NZ_CP148089.1 | GCF_037482275.1 | Barran et al., 2001 |
| *Sinorhizobium meliloti* 2011 | NC_020527.1 | GCF_000346065.1 | Sallet et al., 2013 |
| *Sinorhizobium meliloti* AK21 | NZ_CP026526.1 | GCF_009664245.1 | |
| *Sinorhizobium meliloti* AK76 | NZ_CP066361.1 | GCF_016406285.1 | |
| *Sinorhizobium meliloti* AK83 | NC_015591.1 | GCF_000147795.2 | Galardini et al., 2013 |
| *Sinorhizobium meliloti* B399 | NZ_CP019486.1 | GCF_002302375.1 | |
| *Sinorhizobium meliloti* B401 | NZ_CP019483.1 | GCF_002302355.1 | |
| *Sinorhizobium meliloti* BIM B-442D | NZ_CP123005.1 | GCF_029854455.1 | |
| *Sinorhizobium meliloti* BL225C | NC_017324.1 | GCF_000147775.2 | Galardini et al., 2013 |
| *Sinorhizobium meliloti* CCMM B554 (FSM-MA) | NZ_CP019585.1 | GCF_002215195.1 | |
| *Sinorhizobium meliloti* GR4 | NC_019848.2 | GCF_000320385.2 | |
| *Sinorhizobium meliloti* HM006 | NZ_CP021830.1 | GCF_002197165.1 | |
| *Sinorhizobium meliloti* KH35c | NZ_CP021827.1 | GCF_002197105.1 | |
| *Sinorhizobium meliloti* KH46 | NZ_CP021824.1 | GCF_002197465.1 | |
| *Sinorhizobium meliloti* L6-AK89 | NZ_CP085527.1 | GCF_020684825.1 | |
| *Sinorhizobium meliloti* LPU88 | NZ_CP135242.1 | GCF_042980585.1 | |
| *Sinorhizobium meliloti* MABNR56 | NZ_CP146207.1 | GCF_037023865.1 | |
| *Sinorhizobium meliloti* Rm41 | NZ_CP021809.1 | GCF_002197045.1 | |
| *Sinorhizobium meliloti* RMO17 | NZ_CP009145.1 | GCF_000747295.1 | Toro et al., 2014 |
| *Sinorhizobium meliloti* RRI128 | NZ_CP088115.1 | GCF_021052665.1 | |
| *Sinorhizobium meliloti* RU11/001 | NZ_CP021217.1 | GCF_001050915.2 | |

| Strain | Accession Number | Assembly Accession | Reference |
|---|---|---|---|
| *Sinorhizobium meliloti* S35m | NZ_CP065021.1 | GCF_015689095.1 | |
| *Sinorhizobium meliloti* SM11 | NC_017327.1 | GCF_000218265.1 | Schneiker-Bekel et al., 2011 |
| *Sinorhizobium meliloti* T073 | NZ_CP021805.1 | GCF_002197145.1 | |
| *Sinorhizobium meliloti* USDA1021 | NZ_CP021801.1 | GCF_002197445.1 | |
| *Sinorhizobium meliloti* USDA1106 | NZ_CP021798.1 | GCF_002197065.1 | |
| *Sinorhizobium meliloti* USDA1157 | NZ_CP021794.1 | GCF_002197025.1 | |
